# Brain functional connectivity as a mediator between hematological metrics and cognitive decline in children with beta-thalassemia major

**DOI:** 10.1101/2024.11.06.622267

**Authors:** Shumin Xu, Jiazhen Wu, Yaowen Li, Xinyi Liu, Sixi Liu, Xiaodong Wang, Gui Huang, Mengting Liu, Hongwu Zeng

**Affiliations:** Department of Radiology, Shenzhen Children’s Hospital, Shenzhen, China; Department of Electronic Engineering, Shantou University, Shantou, 515063, Guangdong, China; Department of Radiology, Jiangmen Central Hospital, Jiangmen, China; School of Biomedical Engineering, Sun Yat-sen University, Shenzhen, 518107, China

**Keywords:** beta-thalassemia major, cognitive impairment, fMRI, brain network, connectome-based predictive modeling, hematological abnormalities

## Abstract

Children with beta-thalassemia major (β -TM) are at risk of cognitive impairment, particularly in learning and memory. While cognitive deficits in the general population have been extensively studied, brain-based predictors for β-TM children remain unexplored. This study used connectome-based predictive modeling (CPM) to examine whole-brain functional connectivity in 60 participants to predict cognitive performance. β-TM children showed impaired cognitive abilities, as evidenced by lower Wechsler Intelligence Scale scores compared to controls. The identified brain regions, though not typically linked to cognitive performances, exhibited functional connectivity patterns associated with β-TM, as confirmed by network-based statistics. Correlations between task performance, functional connectivity identified by the CPM, and hematological metrics were also examined. Significant correlations were found between the strength of functional network identified by CPM and hematological metrics, particularly hemoglobin levels and red blood cell distribution width. The study demonstrates that hematological factors influence cognitive performance indirectly through specific functional connectivity, providing new insights into the neural underpinnings of cognitive deficits in children with beta-thalassemia major.

## Introduction

Beta-thalassemia major (β-TM below), one of the most severe types of hereditary hemolytic anemia, has a global carrier rate of 1.5% ^1^. The primary treatment for β-TM requires frequent blood transfusions, which over time leads to iron accumulation in the blood, potentially damaging organs such as the heart, liver, and pancreas, and even causing brain damage ^2,3^. A landmark article published in 2019 recommended that all physicians consider cognitive impairment when treating β-TM ^4^. Other studies have also indicated that patients with β-TM might experience brain iron overload due to long-term transfusions, ineffective erythropoiesis, and increased intestinal iron absorption, leading to cognitive impairment ^5-7^. Most affected β-TM patients experience cognitive function impairments, including difficulties in speech, executive function, working memory, and spatial perception ^4,8^. It is noteworthy that the primary concern in patients with beta-thalassemia who require medical attention are school-aged children, where learning and memory impairments are significant issues in this population ^7^.

The neurological effects of β -TM, including cognitive deficits, remain an under-researched area, despite evidence suggesting their significance ^5^. This underscores the need to consider the central nervous system (CNS) in β-TM diagnosis. Currently, the Wechsler Intelligence Scale for Children (WISC) is commonly used in clinical assessments of cognitive function in children with beta-thalassemia major ^9^. While the WISC can directly reflect a child’s cognitive abilities and provides comparable results for clinical diagnosis, it has significant drawbacks. These include a greater focus on phenotypic manifestations of cognitive function, assessment lag, and the provision of only static results, without accurately pinpointing the relevant brain functional areas ^10^. CNS involvement in β-TM has been overlooked, likely because it often presents sub-clinically, without obvious symptoms or significant impact on daily activities. However, several studies have reported thalassemia-related brain damage, such as silent ischemic lesions, thromboembolic events, white matter abnormalities, intracranial artery stenosis, aneurysms, and sensory evoked potential irregularities ^6,11-16^. These abnormalities may contribute to cognitive and neuropsychological dysfunction in β-TM patients.

Other than structural alterations, functional Magnetic Resonance Imaging (fMRI) can provide comprehensive functional information ^17-20^, helping to understand the neural mechanisms underlying cognitive impairments in β-TM patients ^21^ This complements the limitations of scale assessments and structural observations, explores the mechanisms of cognitive damage, enables early identification of cognitive impairments, and offers opportunities for early intervention.

Although a significant body of neuroimaging research has revealed key elements of the neural circuitry involved in cognitive deficit, the understanding of how functional interactions across multiple large-scale networks in the connectome (i.e., functional connectivity throughout the entire brain) relate to cognitive tasks performance in β-TM, especially children with B-TM remains limited. Connectome modeling approaches, which utilize machine learning algorithms and include a cross-validation step ^22,23^, have increasingly been used to predict behavioral phenotypes from large-scale functional networks ^24-26^. These data-driven methods show promise in identifying disruptions in functional interactions among specific networks in the human connectome associated with cognitive tasks performance, potentially aiding in the discovery of brain–behavior associations and biomarkers ^27,28^. However, no study has yet examined whether the functional organization of the brain can predict cognitive deficit in children with B-TM. This study is the first to employ a connectome predictive modeling approach to identify neural markers of childhood β-TM.

This study aimed to identify network connectivity predictive of cognitive task behaviors in β-TM patients using CPM. Additionally, we analyzed the resulting predictive networks concerning hematological metrics to determine if they were influenced by hematological pathologies. We hypothesized that the mechanisms underlying cognitive deficits in β-TM patients might differ from those in healthy controls. Moreover, we anticipated that a data-driven approach could offer novel insights into how cognitive abilities in β-TM patients are impaired and their relationship with hematological parameters.

## Materials and Methods

### 1 Subjects

This study protocol received approval from the ethics committees of our institution, and written informed consent was obtained from all participants. From May 2020 to October 2023, we recruited a total of 27 patients with beta-thalassemia and 38 healthy controls from our institutional referral centers for beta-thalassemia. Two β-TM patients and three healthy subjects were excluded due to image artifacts (n = 3) and significant registration errors (n = 2). The final analysis included 25 B-TM patients (mean hemoglobin level: 3.71 ± 0.51 g/L; distribution width: 42.06 ± 9.21 g/L; mean ferritin level: 36.3 ± 8.79 ng/mL), all recruited from the Pediatric Hematology Unit at Shenzhen Children’s Hospital.

Exclusion criteria for participants were: (1) presence of other mental disorders, personality disorders, or psychotropic drug dependence; (2) inability to cooperate during MRI examination, resulting in poor image quality; (3) history of other organic or metabolic brain diseases; (4) other MRI contraindications.

To minimize confounding factors related to cerebrovascular involvement, such as environmental and dietary influences, healthy control participants were primarily recruited from the relatives or acquaintances of the patients. None of the control subjects were recruited from a hospital setting, and they were subjected to the same exclusion criteria as the patient group. Prior to undergoing MRI, subjects participated in an interview to screen out those with major cerebrovascular risk factors, such as hypertension or diabetes, ensuring their exclusion from the study.

### 2 Clinical cognitive function assessment

We conducted behavioral tests on both healthy children and those with illnesses, using the latest version (Edition IV) of the Wechsler Intelligence Scale (WIS). This test covers four areas: verbal comprehension, perceptual reasoning, working memory, and processing speed. For our analysis, the WIS scores are calculated as the sum of the scores from these four areas.

### 2 Image acquisition and preprocessing

Structural brain imaging data were obtained on a 3-Tesla MRI unit (Magnetom Skyra, Siemens Medical Solutions, Erlangen, Germany) with a 20-channel head coil array, located at Shenzhen Children’s Hospital. A T1-weighted three-dimensional magnetization prepared-rapid gradient echo (3D-MPRAGE) sequence was acquired with the following parameters: 176 contiguous sagittal slices, 1 mm×1 mm×1 mm raw voxel size, repetition time (TR) =2300 ms, echo time (TE) =2.26 ms, field of view (FOV) =256 mm, number of averages =1, slice oversampling =18.2%. Besides, a T2-weighted sequence with TR=2300 ms and TE=10.6 ms was performed before T1-weighted scanning to exclude organic cerebral lesions. fMRI data was obtained by using an echo-planar imaging (EPI) sequence with the following parameters: repetition time = 2000ms, echo time = 30ms, flip angle = 90 degrees, slice thickness/gap = 4.0/0.0 mm, matrix = 64*64, field of view = 24*24 cm2, 32 axial slices. 130 volumes.

All participants were asked to lie still and stay awake with their eyes closed during scanning. A surveillance camera was set to monitor the subjects for fear of accidents. Foam pads were used to restrain head movements and earplugs were added to drown the scanner noise. Once motion artifacts presented, the subject was requested to receive MRI rescanning immediately to ensure better image quality.

### 3 Imaging preprocessing

Functional imaging data were processed using SPM12 software (https://www.fil.ion.ucl.ac.uk/spm/). For each participant, the initial 10 volumes were discarded to achieve dynamic equilibrium and adaptation to the scanning conditions. The remaining functional images were then adjusted for slice timing to correct time delays between slices. Head motion was corrected using a six-parameter rigid body transformation during the realignment analysis. Participants with head motion exceeding 3 mm in translation or 1.5° in rotation were excluded from further analysis; however, no participants met these exclusion criteria in this study. Each participant’s structural image was coregistered to the mean functional image and segmented into gray matter, white matter (WM), and cerebrospinal fluid (CSF) for normalization. All functional images were spatially normalized to the standard Montreal Neurological Institute (MNI) space with an isotropic voxel size of 3 mm. To enhance the signal-to-noise ratio, the normalized functional images were smoothed using a 4 mm full-width at half-maximum Gaussian filter. A linear detrending and temporal bandpass filtering (0.01 Hz to 0.08 Hz) procedure was applied to mitigate the effects of low-frequency drift and high-frequency physiological noise (Biswal et al., 1995). Additionally, several sources of spurious variance, along with their temporal derivatives, were addressed using linear regression to reduce physiological noise and remove artifacts. This included averaged signals from WM, CSF, and six head motion parameters.

### 4 Functional connectivity construction

To construct the whole-brain functional connectivity (FC) matrix, the Anatomical Automatic Labeling (AAL) template was employed to segment the brain into 116 regions of interest (ROIs), comprising 78 cortical, 12 subcortical, and 26 cerebellar regions (Tzourio-Mazoyer et al., 2002). For each ROI, a representative time series was obtained by averaging the time series of all voxels within the region. Pearson’s correlation analysis was then conducted between each pair of ROIs to determine the correlation coefficients, which were subsequently normalized using a Fisher z-score transformation. This process resulted in a symmetric functional connectivity matrix (116 × 116) for each participant. The upper triangular portion of the adjacency matrix was extracted and converted into a vectorized feature space consisting of 6670 dimensions.

### 5 Network based statistics (NBS)

To identify specific pairs of brain regions with altered functional connectivity in cirrhotic patients, we used the network-based statistic (NBS) approach ^29^. First, we conducted two-sample one-tailed t-tests on each connection that was significantly nonzero (p < 0.05, Bonferroni corrected) in at least one participant. A primary threshold (p < 1e-4 in this study) ^30,31^ was then applied to define a set of suprathreshold links, within which connected components and their sizes (number of links in these components) were determined.

To estimate the significance of each component, a null distribution of connected component size was empirically derived using a nonparametric permutation approach with 10,000 permutations. For each permutation, subjects were randomly reassigned into two groups, and two-sample one-tailed t-tests were performed on the same set of connections. The same primary threshold (p < 1e-4) was used to generate suprathreshold links, and the size of the maximal connected component was recorded. The corrected p-value for a connected component of size M (number of edges) found in the right grouping of control subjects and patients was determined by calculating the proportion of the 10,000 permutations where the maximal connected component size was larger than M.

NBS was applied to compare functional connectivity (FC) differences between control subjects and β-TM patients, revealing the disrupted connectivity induced by β-TM.

### 6 Connectome-based predictive modeling (CPM)

The Connectome-based Predictive Modeling (CPM) was conducted to predict WIS scores using MATLAB scripts. CPM employs connectivity matrices and WIS scores from individuals to create a predictive model for behavioral data based on connectivity matrices. In this process, edges and behavioral data from the training dataset are correlated using regression analyses (Pearson’s correlation or partial correlation when controlling for other behavioral variables or covariates) to identify positive and negative predictive networks. Positive networks show increased edge weights (connectivity) associated with the variable of interest, while negative networks show decreased edge weights associated with the variable.

Single-subject summary statistics are generated by summing the significant edge weights in each network, which are then used in predictive models assuming linear relationships with behavioral data. The predictive networks and summary score model identified from the training data are applied to the test dataset to predict behavior.

Leave-one-out cross-validation was performed to determine if cognitive score could be predicted based on the connectivity profile of an unseen individual. In this approach, the predicted value for a single subject (the “left-out” participant) is generated using data from all other participants (N-1) as the training dataset. This iterative process involves setting aside one subject’s data as the test set while using the remaining N-1 subjects’ data as the training set, until all subjects have a predicted value. Each iteration includes: (i) feature selection, identifying edges with a significant relationship to aggression severity in the training set and classifying them based on their sign (positive or negative); (ii) model building, fitting linear regressions between aggression and connectivity strength in positive- and negative-feature networks using the training data; and (iii) prediction, inputting data from the excluded subject into each model to generate a predicted aggression score ^32,33^.

After all iterations, CPM model performance was assessed by correlating the predicted and observed WIS scores.

### 7 Localization of predictive networks

Predictive networks were summarized at various levels of data reduction, including edge, node, and network levels ^23^. The overlap of nodes with macroscale brain regions (such as the motor cortex and cerebellum) was determined based on anatomical labels from Finn et al. ^24^. The overlap of nodes with canonical functional network localizations (like frontoparietal and sensorimotor networks) was based on the functional networks from Nobel et al. ^34^. Additionally, for each node, the network theory measure “degree” was calculated as the sum of the number of edges for each node within the predictive networks.

Visualizations of predictive edges were created using BioImage Suite Web (https://bioimagesuiteweb.github.io/alphaapp/index.hβ-TMl) ^35^. High-degree nodes were defined as the top nodes with the most edges or connections across all iterations of the predictive model.

### 8 Significance of CPM performance

For connectome analyses, the correspondence between predicted and actual values, or model performance, was assessed using Pearson’s correlation. Negative correlations were set to zero. During cross-validation, analyses in the left-out folds are not independent, leading to an overestimation of degrees of freedom for parametric p-values. Therefore, permutation testing was performed. To generate null distributions for significance testing, we randomly shuffled the correspondence between behavioral variables and connectivity matrices by permuting subject assignments for behavioral variables 1000 times and reran the CPM analysis with the shuffled data. Based on these null distributions, the p-values for predictions were calculated ^36^. Given our hypothesis of a positive association between predicted and actual values, one-tailed p-values are reported.

### 9 Follow-up analyses

To determine the construct specificity of high-degree nodes in predicting WIS scores, follow-up tests were carried out by retaining the high-degree nodes and all edges connected to them (i.e., removing all other edges). This approach aimed to assess the robustness of these networks in predicting WIS scores.

## Results

### 1 Demographic information and clinical variables

In total, 25 individuals with β-TM were included in the study, consisting of 15 males with an average age of 9.83 years (standard deviation ± 2.09 years). For normal control group, 35 individuals were included in the study, consisting of 19 males with an average age of 9.72 years (standard deviation ± 1.81 years). Comprehensive demographic and clinical details of these groups are provided in Table 1.

**Table 1.**
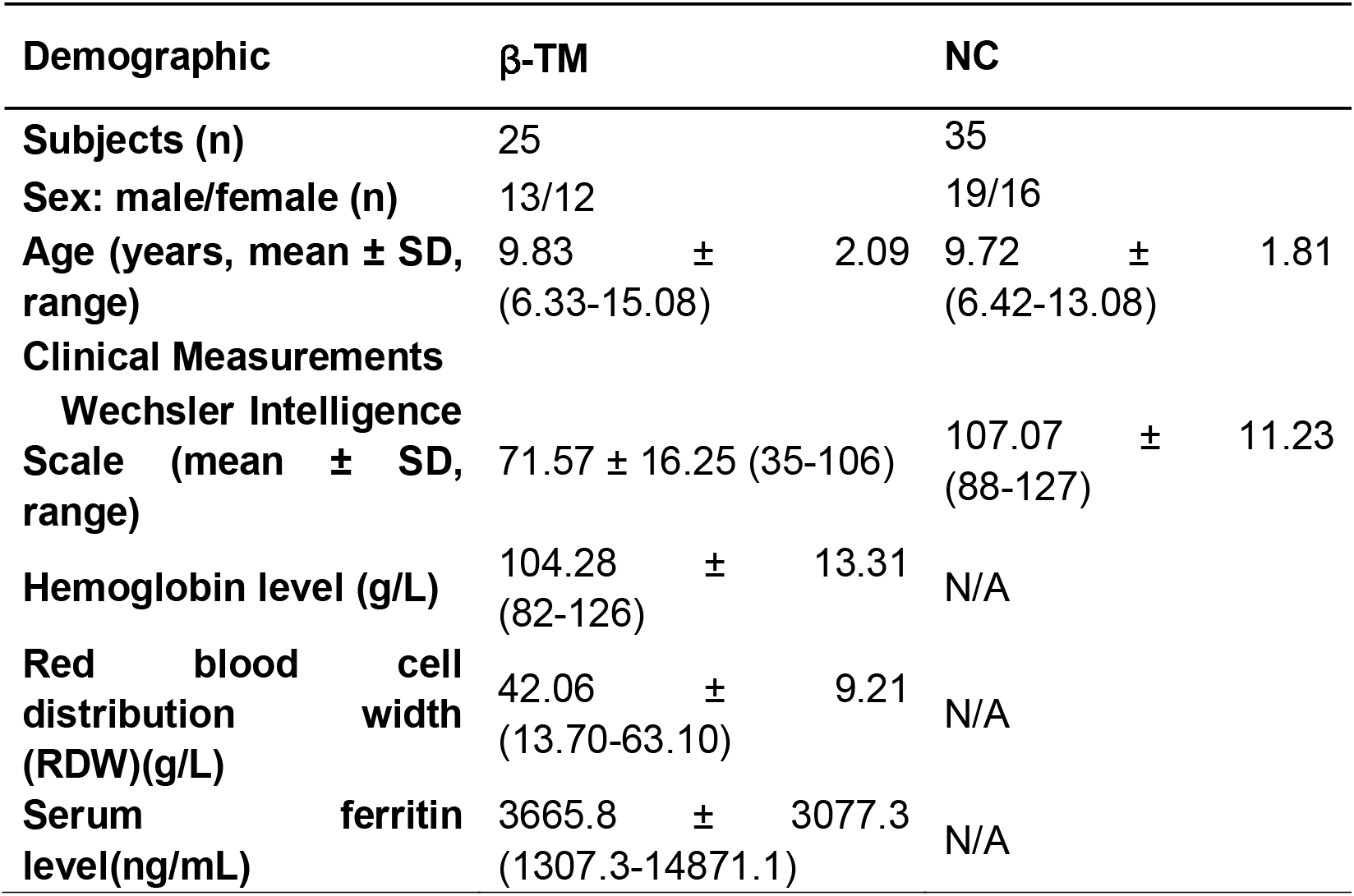
Demographic and clinical characteristics for B-TM and NC subjects

### 2 Between group differences in clinical measurements

Overall, individuals with β-TM exhibited significantly lower WIS scores (p < 0.001) compared to normal controls, suggesting impaired cognitive function in β-TM patients. Detailed information on the WIS scores for these groups is presented in Table 1.

Hematological metrics such as hemoglobin level, red blood cell distribution width (RDW), and serum ferritin level were exclusively measured in B-TM patients in this study. In β-TM patients, WIS scores showed a negative correlation with serum iron levels (p’s < 0.001).

### 3 Disrupted functional connectivity in β-TM

NBS was initially applied to compare functional connectivity (FC) differences between control subjects and β-TM patients. Using a cluster-defining threshold of p < 1e-4 (as explained in Materials and Methods), a single network consisting of 72 connections among 41 brain regions was identified, showing decreased functional connectivity in the B-TM group (p = 0.012). This decreased connectivity was primarily found among cerebellum’s posterior lobules, including Crus II–VIIb, cerebellum X, cortical regions in motor ares such as Heschl’s gyrus (HG) and the superior temporal gyrus (STG), et al. (Fig. 1). The top 5 regions found with highest degrees are: right Cerebellum XIIb, right cerebellum XIII, left and right cerebellum Curs II, and right superior temporal cortex.

**Fig. 1.**
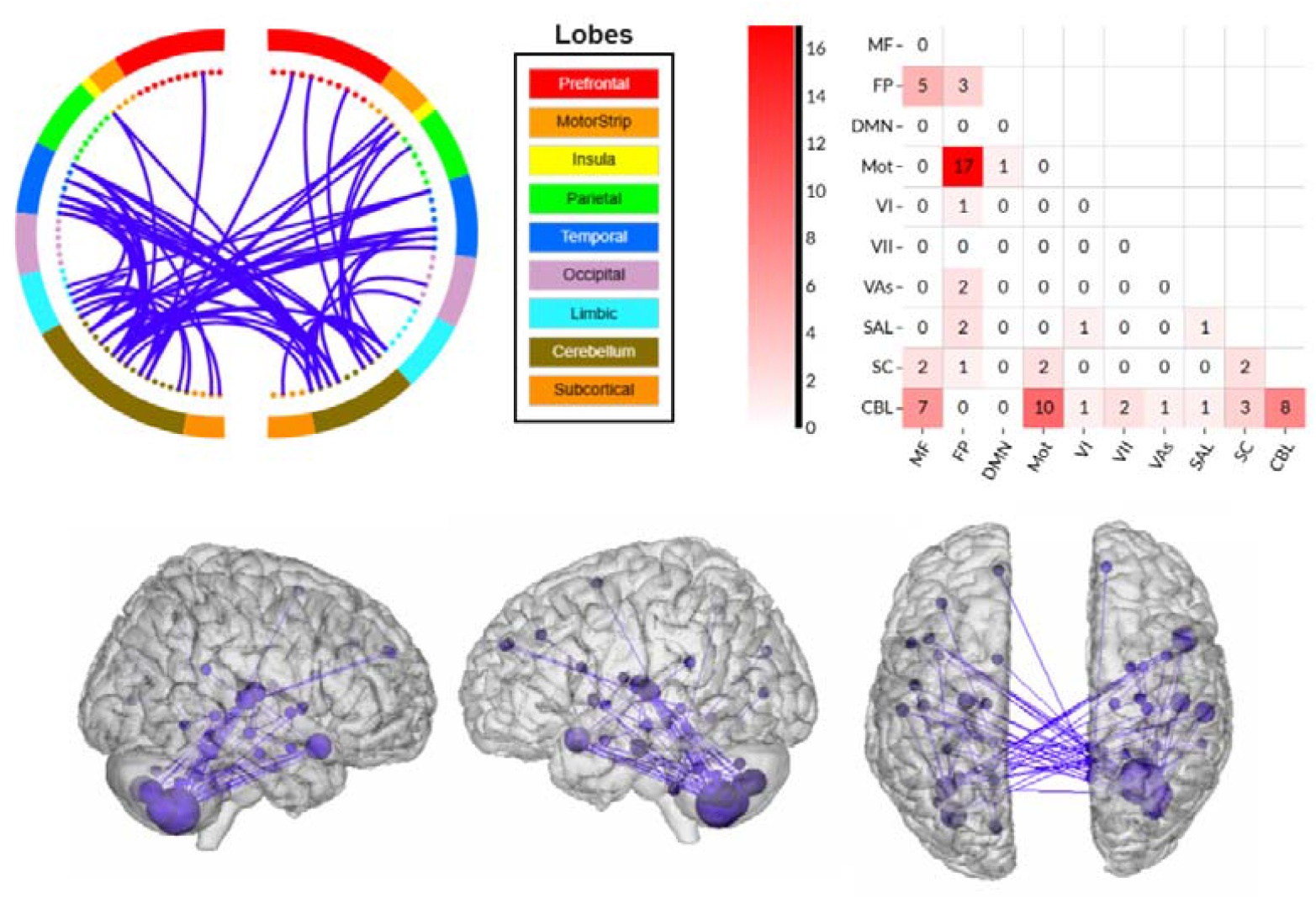
Brain-wide functional connectivity impairment in B-TM. B-TM patients exhibit a large number of decreased/impaired functional connectivity across wide range of the brain regions (blue edges). No increased functional connectivity was found in B-TM patients. Larger spheres indicate nodes with more edges, and smaller spheres indicate nodes with fewer edges. Within- and between-network connectivity for the impaired functional networks are summarized using canonical networks. Cells represent the total number of edges connecting nodes within and between each network, with darker colors indicating a greater number of edges.

### 4 Prediction of WIS score

The comprehensive CPM model demonstrated that brain-wide connectivity patterns could predict cognitive task performance, with only positive networks identified (r = 0.30, p = 0.016 via permutation testing) (Fig. 2A). Follow-up comparisons were performed to assess the impact of potential covariates on the CPM model’s prediction of WIS scores. Models accounting for potential covariates also successfully predicted WIS scores, showing similar predictive performance when controlling for age (r = 0.27, p = 0.021) and sex (r = 0.28, p = 0.019). Further analyses were conducted using ten-fold cross-validation, yielding similar results; however, as anticipated, the correlation coefficient was slightly lower with ten-fold cross-validation compared to leave-one-out cross-validation (r = 0.25, p = 0.031).

**Fig. 2.**
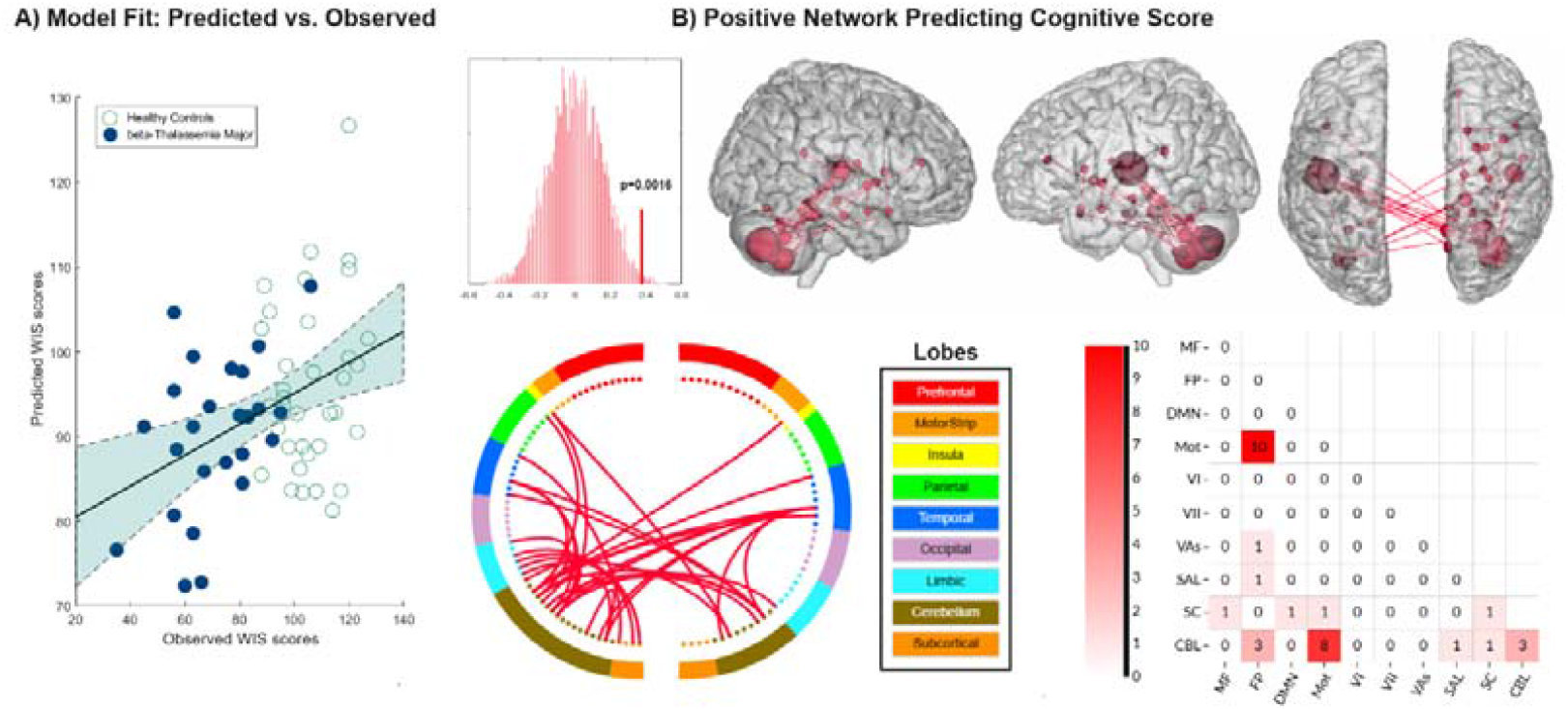
Brain-wide functional connectivity predicts cognitive task performance. A) The correspondence between observed (x-axis) and predicted (y-axis) WIS scores generated using CPM. Despite the clinical complexity of the population, CPM successfully predicted WIS scores (p=0.0016, permutation testing). Predictions remained significant in follow-up analyses controlling for covariates including age, and sex. Histogram shows distribution of Pearson correlation (r) values from 5000 iterations of randomly shuffled ratings of aggression severity used to nonparametrically determine p values. B) Positive (red) predicting aggression. For positive networks, increased edge weights (i.e., increased functional connectivity) predict WIS scores. No negative network was found in current study. Larger spheres indicate nodes with more edges, and smaller spheres indicate nodes with fewer edges.

**Fig. 3.**
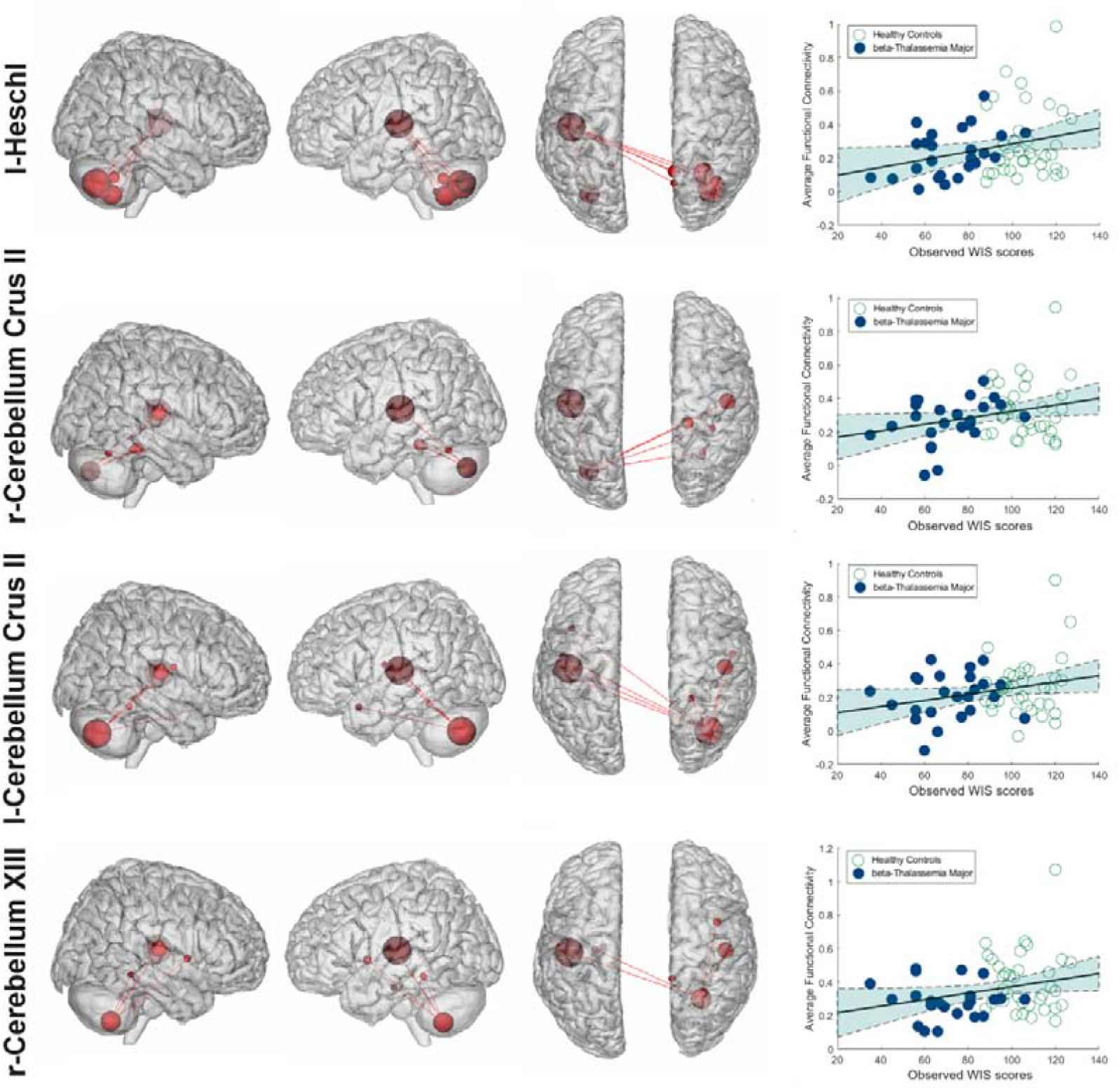
Follow up analyses tested the sensitivity of high-degree nodes (i.e., most predictive features) in predicting WIS scores. Regions emerging as highly predictive of aggression in the connectome model include the r-Cerebellum XIII, l-Cerebellum Crus II, r-Cerebellum Crus II, and –Heschl. We preserved the high-degree nodes from CPM and all their connected edges, while removing all other edges.

### 5 Network anatomy and localization of circuits

Fig. 2B provides a detailed overview of the cognitive networks, where only positive networks were detected, with no connectivity observed in negative networks. The highest-degree nodes (those with the most connections) in the positive network were predominantly located in the posterior lobules of the cerebellum, including Crus II–VIIb, cerebellum VIII, as well as cortical regions in motor areas such as Heschl’s gyrus (HG) and the superior temporal gyrus (STG), among others. These findings are notably similar to those shown by Network-Based Statistic (NBS) analysis. The top four regions with the highest degree of connectivity were the left Heschl cortices, left and right cerebellum XIIb, and right cerebellum XIII. On a network level, the overall between-network connectivity was primarily characterized by interactions between the frontoparietal and motor networks, as well as connections among the cerebellum and motor networks.

### 6 Follow-up analyses

Follow-up analyses examined the sensitivity of high-degree nodes (i.e., most predictive features) in forecasting WIS scores. We preserved the high-degree nodes from CPM and all their connected edges, while removing all other edges. Results revealed that after FDR correction, top four high-degree nodes from CPM network still show significant predictive power for WIS score (r = 0.24,p = 0.039 for left Heschl cortices, r = 0.24, p = 0.041 for right cerebellum XIIb, r = 0.23, p = 0.046 for right cerebellum XIIb, and r = 0.22, p = 0.048 for right cerebellum XIII.)

### 7 Predictive power between β-B-TM and controls

To determine whether the identified network serves as a generalized predictor of cognitive task performance for all participants or specifically indicates cognitive deficits in β-TM patients, we conducted separate regression analyses using the network strength with WIS scores for both β-TM and control groups. The results indicated that the network was predictive only for β-TM children (r = 0.30, p = 0.031; Fig. 4A), and not for the control group. Further analyses revealed that the node degree for the left Heschl’s gyrus (r = 0.29, p = 0.035) and the right Cerebellum Crus II (r = 0.27, p = 0.042) were significant predictors of WIS scores in β-TM children. However, none of the node degrees in these regions predicted WIS scores in healthy controls.

**Fig. 4.**
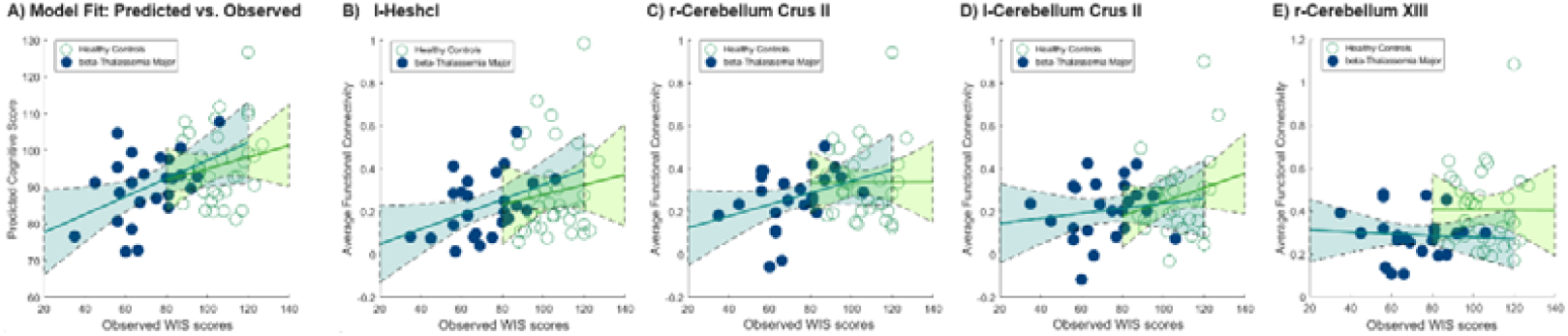
Separate regression analyses were performed for the β-TM and control groups independently. Significant relationships with cognitive task performance were identified exclusively in β-TM patients, regardless of the networks identified in CPM or the sensitivity of high-degree nodes.

### 8 Association of task performance, functional network strength and hematological metrics

Univariate analyses showed that higher hemoglobin levels (r = 0.32; corrected p = 0.017, Fig. 5A) and lower red blood cell distribution width (r = 0.35; corrected p = 0.009, Fig. 5B) were associated with increased functional connectivity as identified by CPM.

**Fig. 5.**
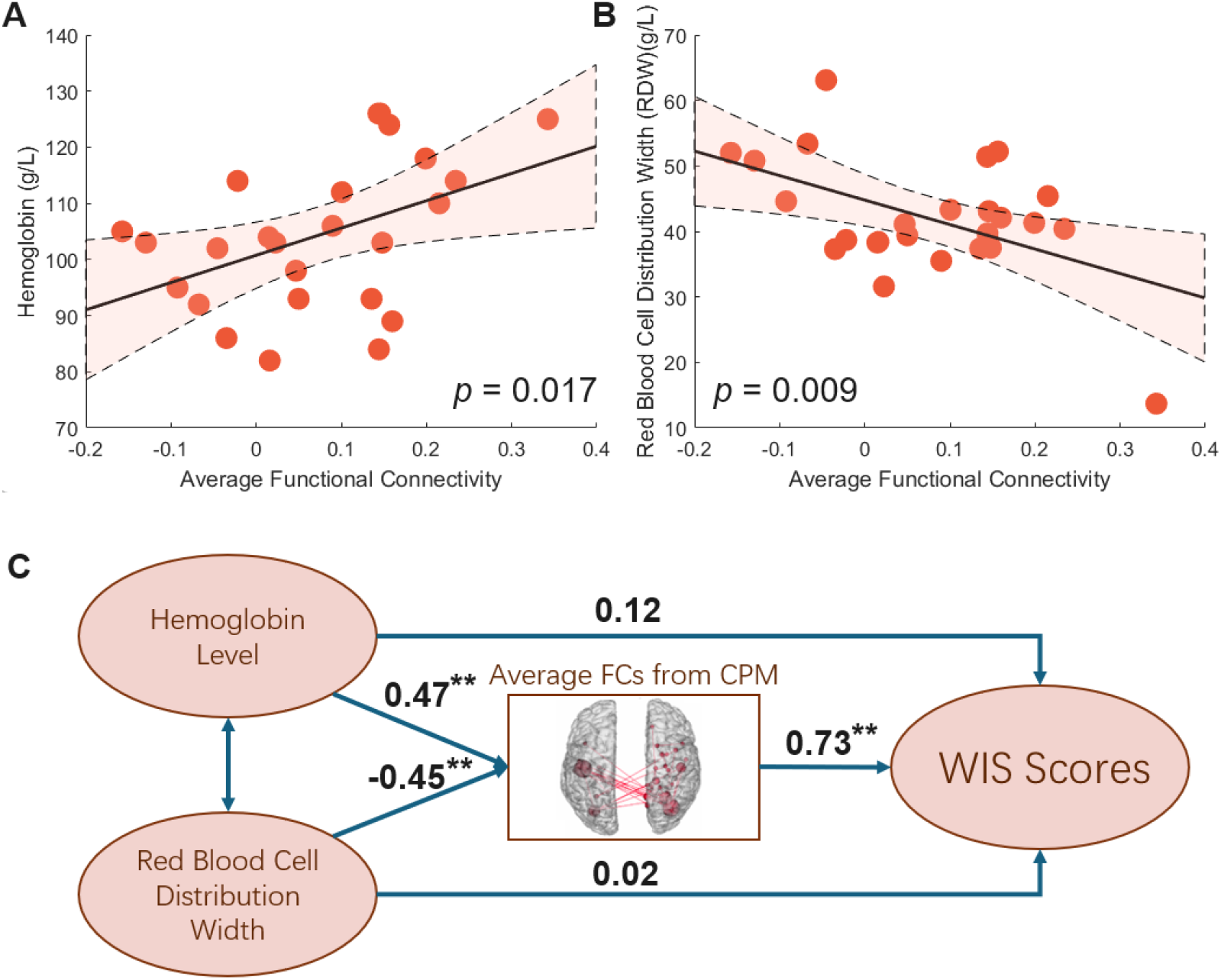
Univariate analyses revealed that higher A) hemoglobin levels and B) lower red blood cell distribution width was linked to increased functional connectivity, as identified by CPM. C) SEM suggests that the functional connectivity observed in CPM mediates the impact of hematological abnormalities on WIS scores.

The structural equation model (SEM), depicted in Fig. 5C with standardized estimates, indicated a significantly acceptable model fit based on model-fit statistics (p < 0.001). The average functional connectivity from the CPM network mediated the relationship between both hemoglobin levels (p = 0.0021, Fig. 5C) and red blood cell distribution width (p = 0.0024; Fig. 5C) and WIS scores. Importantly, the direct relationships between the hematological metrics and WIS scores were not significant, underscoring the mediating role of brain functional connectivity in the cognitive deficit pathway for β-TM patients.

## Discussion

In this study, we investigated the impairment of functional connectivity in β-TM children using comprehensive whole-brain functional connectivity comparisons and connectome-behavior predictive mapping approaches. Our findings revealed that the impaired functional connectivity in β-TM may influences cognitive performance.

### Cognitive and functional impairments in β-TM

We observed a notable reduction in cognitive abilities in β-TM children, as measured by the Wechsler Intelligence Scales. Children with β-TM are susceptible to excessive iron accumulation in the brain, resulting from chronic blood transfusions, ineffective erythropoiesis, and increased intestinal iron absorption. This accumulation may contribute to cognitive deficits ^3^.

In the imaging studies, notable findings emerged from the NBS-based comparison and post-hoc analysis among groups. β-TM patients displayed widespread disruptions in functional connectivity compared to control subjects, indicating more severe brain impairment due to β-TM. Notably, no significant increases in functional connectivity were observed in β-TM children. The disrupted functional connectivity primarily occurred between the motor network and fronto-parietal networks, as well as between the cerebellum and fronto-parietal networks, the cerebellum and motor networks, and within the cerebellum network itself.

It’s worth noting that the frontoparietal network here we are not indicating the cortices in frontal and parietal lobes, Whereas, we were suggesting the an important part of the cerebellum, the cerebellum’s posterior lobules, including Crus II–VIIb ^34,37^, which are thought correlated with the frontoparietal network (FPN), and was always thought be part of the frontoparietal network. It is involved in a number of non-motor, cognition and executive functions. These functions include working memory, planning, organizing, and strategy formation, which are important for creative divergent thinking ^38^.

When referring to the motor cortex, we primarily mean the Heschl cortex and the superior temporal cortices. Both Heschl’s gyrus (HG) and the superior temporal gyrus (STG) are crucial for auditory processing and language ^39^. Heschl’s gyrus is also known as the transverse temporal gyrus, is a significant component of the posterior portion of the STG and serves as the primary auditory area. It is involved in various functions, including speech production, phonologic retrieval, semantic processing, and language comprehension ^40^. Superior Temporal Gyrus (STG) is is divided into anterior, middle, and posterior parts, encompassing HG and Wernicke’s area. Bilateral activation of the STG enables the perception of speech sounds of varying lengths, such as syllables, words, and pseudowords ^41^. Damage to the STG on both sides can result in pure word deafness, highlighting its role in mapping acoustic signals to abstract speech. Additionally, the STG is essential for short-term auditory sensory memory.

Functional connectivity between these regions facilitates the integration of perceptual information with cognitive functions. Disruptions in this connectivity can result in difficulties processing and understanding perceptual information, thereby impairing overall cognitive performance.

Recent anatomical and functional studies have highlighted that various regions of the cerebellum are involved in a broad spectrum of cognitive functions beyond its traditional role in sensorimotor control ^42,43^. The cerebellum has been implicated in executive function ^44^, attention ^45^, and emotional processing ^46^. Specifically, cerebellum lobe X, a crucial hub in the PMN, has been identified as a non-motor area in recent research ^47^, and is thought to be associated with visual working memory and visual recognition ^42^. The exact role of the cerebellum in cognitive performance for B-TM patients warrants further investigation in future studies.

### Functional decoding of WIS scores in β-TM

Wechsler Intelligence Scale includes four aspects: Verbal Comprehension Index, Perceptual Reasoning Index, Working Memory Index, and Processing Speed Index, which is a comprehensive evaluatation of the cognitive functions of children with severe β -TM and healthy control children. The functional connectivity found from CPM indicating the cognitive function is largely overlap with the impaired functional connectivity obtained from NBS, implying that the impaired FCs actually impaired the cognitive ability.

Techniques such as the transverse relaxation rate ^48^, susceptibility-weighted imaging (SWI) ^49^, and quantitative susceptibility mapping (QSM) ^14^ in brain magnetic resonance imaging (MRI) can be used for in vivo and quantitative assessments of cerebral iron levels. Previous studies using these methods have reported iron overload and deposition in the motor, temporal, and sub-cortical gray matter areas ^3,50,51^, such as cerebellum. These findings are consistent with the impaired hub in functional connectivity identified in our study, further supporting the hypothesis of iron overload in these areas. Future research could include more direct investigations of iron content overload using modalities like QSM and SWI to substantiate these findings.

### Functional connectivity mediates hematological abnormalities and cognitive decline

This study uses structural equation modeling (SEM) to demonstrate the causal effects of hematological factors on cognitive impairments in β-TM patients ^52^. Specifically, brain connectivity identified by CMP during the scan mediated the relationship between hematological metric scores, such as hemoglobin levels and red blood cell distribution width, and WIS scores (Fig. 5). This may suggest that hematological abnormalities in β -TM patients negatively impact cognitive function through alterations in the brain’s functional architecture. However, SEM did not find significant direct evidence of a relationship between hematological metric scores and WIS scores. The underlying pathophysiology through which these hematological pathologies impair cognitive abilities likely involves both external and internal factors, including excessive brain iron accumulation from chronic blood transfusions, ineffective erythropoiesis, and increased intestinal iron absorption, all potentially leading to cognitive impairments.

### Data-driven approach may find novel and robust neuromarkers

The data-driven approach based on Network-Based Statistics (NBS) and Connectome-Based Predictive Modeling (CPM) employed in this study identified novel connections used for group comparison and predicting cognitive test outcomes. Previous hypothesis-driven approaches ^53,54^ have restricted the selection of regions of interest to brain areas known to be associated with specific cognitive tasks. In contrast, data-driven feature selection methods ^29,55,56^, like the one adopted in this study, do not have such limitations. Our approach revealed that the neural mechanisms underlying cognitive task processing in β-TM children may affected by brain regions that is not ordinary to be the well-known regions for cognitive intensive tasks, which might be focused on frontal and parietal cortices of the brain. This finding suggests that specific, yet novel brain connections may be impaired in the cognitive tasks of interest for patients with β-TM.

Group-wise analysis also reveals that the functional connectivity for cognitive tasks using CPM only predict the WIS scores for β-TM children but not healthy children (Fig. 4). This may suggest that the found network is particularly exsert influence on cognitive ability for β-TM patients.

### Limitations and future directions

This study has several limitations. While we identified functional connectivity detrimental to cognitive performance, the underlying mechanisms remain unclear, particularly how iron accumulation affects functional connectivity. Future studies should consider in vivo and quantitative assessments of cerebral iron levels using techniques like transverse relaxation rate, susceptibility-weighted imaging (SWI), and quantitative susceptibility mapping (QSM) in brain MRI. Due to the scarcity of B-TM data, the findings in this study lack external validation. Additionally, our failure to collect blood samples from the control group limits our ability to compare hematological metrics between β-TM patients and healthy individuals. This oversight hinders our analysis of how differences in blood composition, such as iron levels and red blood cell counts, might distinguish β-TM patients from controls. The absence of hematological metrics from control subjects restricts our understanding of the overall influence of these blood characteristics on cognitive functions. Our data-driven method also has limitations. It is challenging to clarify the role of identified functional networks or brain regions when their original functions are not associated with the target cognitive function. Additionally, the modest sample size may inflate some reported effect sizes, particularly given the high-dimensional feature space, leading to a higher false positive rate for some results ^57^. Future studies should aim to replicate these findings with larger sample sizes.

## Acknowledgements

The authors thank all subjects involved in the study. This study was supported by Excellent Youth Foundation of Zhejiang Scientific Committee LR24F010002, and Key Program of Zhejiang Scientific Committee LZ23F010002.

## Data and Code Availability Statement

Our native MRI images and code will be made available on request from any qualified investigator who provides a practicable proposal, or for the purpose of replicating procedures and results presented in the current study.

## Authors contributions

**S. X**. performed the research, analysed the data and wrote the paper. **J. W**. analysed the data, and wrote the paper; **Y. L**. prepared the software and analysed the data; **X. L**. critically reviewed the manuscript; **S. L**. collected the data; **X. W**. critically reviewed the manuscript; **G. H**. critically reviewed the manuscript; **M. L**. designed the research study, supervised the study and critically reviewed the manuscript; **H. Z**. designed the research study, supervised the study, critically reviewed the manuscript and provided the funding.

## References

1. Galanello R, Origa R. Beta-thalassemia. Orphanet journal of rare diseases. 2010;5:1–15.

2. Berdoukas V, Farmaki K, Carson S, Wood J, Coates T. Treating thalassemia major-related iron overload: the role of deferiprone. Journal of blood medicine. 2012:119–129.

3. Manara R, Ponticorvo S, Tartaglione I, et al. Brain iron content in systemic iron overload: A beta-thalassemia quantitative MRI study. NeuroImage: Clinical. 2019;24:102058.

4. Tartaglione I, Manara R, Caiazza M, et al. Brain functional impairment in beta - thalassaemia: the cognitive profile in Italian neurologically asymptomatic adult patients in comparison to the reported literature. British journal of haematology. 2019;186(4):592–607.

5. Monastero R, Monastero G, Ciaccio C, Padovani A, Camarda R. Cognitive deficits in beta -thalassemia major. Acta Neurologica Scandinavica. 2000;102(3):162–168.

6. Nemtsas P, Arnaoutoglou M, Perifanis V, Koutsouraki E, Orologas A. Neurological complications of beta-thalassemia. Annals of hematology. 2015;94:1261–1265.

7. Raafat N, Safy UE, Khater N, et al. Assessment of cognitive function in children with beta-thalassemia major: a cross-sectional study. Journal of child neurology. 2015;30(4):417–422.

8. Pholngam N, Jamrus P, Viwatpinyo K, et al. Cognitive impairment and hippocampal neuronal damage in β-thalassaemia mice. Scientific Reports. 2024;14(1):10054.

9. Hrabok M, Brooks BL, Fay-McClymont TB, Sherman EM. Wechsler Intelligence Scale for Children-(WISC-IV) short-form validity: A comparison study in pediatric epilepsy. Child Neuropsychology. 2014;20(1):49–59.

10. Hessl D, Nguyen DV, Green C, et al. A solution to limitations of cognitive testing in children with intellectual disabilities: the case of fragile X syndrome. Journal of Neurodevelopmental Disorders. 2009;1:33–45.

11. Karimi M, Khanlari M, Rachmilewitz EA. Cerebrovascular accident in β-thalassemia major (β - TM) and β - thalassemia intermedia (β - TI). American journal of hematology. 2008;83(1):77–79.

12. Musallam KM, Beydoun A, Hourani R, et al. Brain magnetic resonance angiography in splenectomized adults with β-thalassemia intermedia. European journal of haematology. 2011;87(6):539–546.

13. Pazgal I, Inbar E, Cohen M, Shpilberg O, Stark P. High incidence of silent cerebral infarcts in adult patients with beta thalassemia major. Thrombosis research. 2016;144:119–122.

14. Qiu D, Chan G-F, Chu J, et al. MR quantitative susceptibility imaging for the evaluation of iron loading in the brains of patients with β-thalassemia major. American Journal of Neuroradiology. 2014;35(6):1085–1090.

15. Raz S, Koren A, Levin C. Attention, response inhibition and brain event-related potential alterations in adults with beta - thalassaemia major. British journal of haematology. 2019;186(4):580–591.

16. Taher A, Isma’eel H, Mehio G, et al. Prevalence of thromboembolic events among 8,860 patients with thalassaemia major and intermedia in the Mediterranean area and Iran. Thrombosis and haemostasis. 2006;96(10):488–491.

17. Gosein M, Maharaj P, Balkaransingh P, et al. Imaging features of thalassaemia. The British Journal of Radiology. 2019;92(1096):20180658.

18. Gupta S, Patel K. Case series: MRI features in cerebral malaria. Indian Journal of Radiology and Imaging. 2008;18(03):224–226.

19. Kim JA, Leung J, Lerch JP, Kassner A. Reduced cerebrovascular reserve is regionally associated with cortical thickness reductions in children with sickle cell disease. Brain Research. 2016;1642:263–269.

20. Kirk GR, Haynes MR, Palasis S, et al. Regionally specific cortical thinning in children with sickle cell disease. Cerebral Cortex. 2009;19(7):1549–1556.

21. Raz S, Koren A, Dan O, Levin C. Executive function and neural activation in adults with β -thalassemia major: an event-related potentials study. Annals of the New York Academy of Sciences. 2016;1386(1):16–29.

22. Scheinost D, Noble S, Horien C, et al. Ten simple rules for predictive modeling of individual differences in neuroimaging. NeuroImage. 2019;193:35–45.

23. Yip SW, Kiluk B, Scheinost D. Toward addiction prediction: an overview of cross-validated predictive modeling findings and considerations for future neuroimaging research. Biological Psychiatry: Cognitive Neuroscience and Neuroimaging. 2020;5(8):748–758.

24. Finn ES, Shen X, Scheinost D, et al. Functional connectome fingerprinting: identifying individuals using patterns of brain connectivity. Nature neuroscience. 2015;18(11):1664–1671.

25. Ibrahim K, Noble S, He G, et al. Large-scale functional brain networks of maladaptive childhood aggression identified by connectome-based predictive modeling. Molecular psychiatry. 2022;27(2):985–999.

26. Yip SW, Scheinost D, Potenza MN, Carroll KM. Connectome-based prediction of cocaine abstinence. American Journal of Psychiatry. 2019;176(2):156–164.

27. Cheng Y, Shen W, Xu J, et al. Neuromarkers from whole-brain functional connectivity reveal the cognitive recovery scheme for overt hepatic encephalopathy after liver transplantation. Eneuro. 2021;8(4)

28. Liu M, Backer RA, Amey RC, Splan EE, Magerman A, Forbes CE. Context matters: situational stress impedes functional reorganization of intrinsic brain connectivity during problem-solving. Cerebral Cortex. 2021;31(4):2111–2124.

29. Zalesky A, Fornito A, Bullmore ET. Network-based statistic: identifying differences in brain networks. Neuroimage. 2010;53(4):1197–1207.

30. Liu M, Backer RA, Amey RC, Forbes CE. How the brain negotiates divergent executive processing demands: Evidence of network reorganization in fleeting brain states. NeuroImage. 2021;245:118653.

31. Wang J, Zuo X, Dai Z, et al. Disrupted functional brain connectome in individuals at risk for Alzheimer’s disease. Biological psychiatry. 2013;73(5):472–481.

32. Shen X, Finn ES, Scheinost D, et al. Using connectome-based predictive modeling to predict individual behavior from brain connectivity. nature protocols. 2017;12(3):506–518.

33. Wu X, Yang Q, Xu C, et al. Connectome-based predictive modeling of compulsion in obsessive–compulsive disorder. Cerebral Cortex. 2023;33(4):1412–1425.

34. Noble S, Spann MN, Tokoglu F, Shen X, Constable RT, Scheinost D. Influences on the test–retest reliability of functional connectivity MRI and its relationship with behavioral utility. Cerebral cortex. 2017;27(11):5415–5429.

35. Papademetris X, Jackowski MP, Rajeevan N, et al. BioImage Suite: An integrated medical image analysis suite: An update. The insight journal. 2006;2006:209.

36. Lake EM, Finn ES, Noble SM, et al. The functional brain organization of an individual allows prediction of measures of social abilities transdiagnostically in autism and attention-deficit/hyperactivity disorder. Biological psychiatry. 2019;86(4):315–326.

37. Li W, Han T, Qin W, et al. Altered Functional Connectivity of Cognitive-Related Cerebellar Subregions in Well-Recovered Stroke Patients. Neural Plasticity. 2013;2013(1):452439.

38. Prati JM, Pontes-Silva A, Gianlorenço ACL. The cerebellum and its connections to other brain structures involved in motor and non-motor functions: a comprehensive review. Behavioural Brain Research. 2024:114933.

39. Fernández L, Velásquez C, Porrero JAG, de Lucas EM, Martino J. Heschl’s gyrus fiber intersection area: a new insight on the connectivity of the auditory-language hub. Neurosurgical focus. 2020;48(2):E7.

40. Mustroph ML, Zekelman LR, Golby AJ. Probing the tract organization of language: Heschl’s gyrus fiber intersection area. Neurosurgical Focus. 2020;48(2):E8.

41. Friederici AD, Rüschemeyer S-A, Hahne A, Fiebach CJ. The role of left inferior frontal and superior temporal cortex in sentence comprehension: localizing syntactic and semantic processes. Cerebral cortex. 2003;13(2):170–177.

42. King M, Hernandez-Castillo CR, Poldrack RA, Ivry RB, Diedrichsen J. Functional boundaries in the human cerebellum revealed by a multi-domain task battery. Nature neuroscience. 2019;22(8):1371–1378.

43. Strick PL, Dum RP, Fiez JA. Cerebellum and nonmotor function. Annual review of neuroscience. 2009;32(1):413–434.

44. Koziol LF, Budding DE, Chidekel D. From movement to thought: executive function, embodied cognition, and the cerebellum. The Cerebellum. 2012;11(2):505–525.

45. Osaka N, Osaka M, Kondo H, Morishita M, Fukuyama H, Shibasaki H. The neural basis of executive function in working memory: an fMRI study based on individual differences. Neuroimage. 2004;21(2):623–631.

46. Guell X, Gabrieli JD, Schmahmann JD. Triple representation of language, working memory, social and emotion processing in the cerebellum: convergent evidence from task and seed-based resting-state fMRI analyses in a single large cohort. Neuroimage. 2018;172:437–449.

47. Guell X, Schmahmann J. Cerebellar functional anatomy: a didactic summary based on human fMRI evidence. The Cerebellum. 2020;19(1):1–5.

48. Cheung JS, Au WY, Ha SY, et al. Reduced transverse relaxation rate (RR2) for improved sensitivity in monitoring myocardial iron in thalassemia. Journal of Magnetic Resonance Imaging. 2011;33(6):1510–1516.

49. Qiu D, Chan G, Chan Q, Ha S-Y, Khong P-L. Susceptibility weighted imaging (SWI) for the assessment of iron loading in the brain of beta-thalassemia major patients. International Society of ISMRM.; 2010:

50. Metafratzi Z, Argyropoulou M, Kiortsis D, Tsampoulas C, Chaliassos N, Efremidis S. T 2 relaxation rate of basal ganglia and cortex in patients with β-thalassaemia major. The British journal of radiology. 2001;74(881):407–410.

51. Schweitzer AD, Liu T, Gupta A, et al. Quantitative susceptibility mapping of the motor cortex in amyotrophic lateral sclerosis and primary lateral sclerosis. AJR American journal of roentgenology. 2015;204(5):1086.

52. Liu M, Lu M, Kim SY, et al. Brain age predicted using graph convolutional neural network explains neurodevelopmental trajectory in preterm neonates. European Radiology. 2024;34(6):3601–3611.

53. Garcia-Garcia R, Cruz-Gómez ÁJ, Urios A, et al. Learning and memory impairments in patients with minimal hepatic encephalopathy are associated with structural and functional connectivity alterations in hippocampus. Scientific reports. 2018;8(1):9664.

54. Qi R, Zhang L, Wu S, et al. Altered resting-state brain activity at functional MR imaging during the progression of hepatic encephalopathy. Radiology. 2012;264(1):187–195.

55. Liu M, Amey RC, Backer RA, Simon JP, Forbes CE. Behavioral Studies using large-scale brain networks–methods and validations. Frontiers in Human Neuroscience. 2022;16:875201.

56. Rosenberg MD, Finn ES, Scheinost D, et al. A neuromarker of sustained attention from whole-brain functional connectivity. Nature neuroscience. 2016;19(1):165–171.

57. Button KS, Ioannidis JP, Mokrysz C, et al. Power failure: why small sample size undermines the reliability of neuroscience. Nature reviews neuroscience. 2013;14(5):365–376.

